# Specific Wavelengths of Light Modulate Honey Bee Locomotor Activity

**DOI:** 10.1101/2025.10.22.683871

**Authors:** Babur Erdem, Iremnur Fidan, Ali Emre Turgut, Erol Sahin, Ayse Gul Gozen, Hande Alemdar

**Affiliations:** Center for Robotics and Artificial Intelligence (ROMER), Middle East Technical University, Ankara, Türkiye; Department of Biology, Faculty of Arts and Sciences, Düzce University, Düzce, Türkiye; Department of Mechanical Engineering, Middle East Technical University, Ankara, Türkiye; Department of Computer Engineering, Middle East Technical University, Ankara, Türkiye; Department of Biological Sciences, Middle East Technical University, Ankara, Türkiye

**Keywords:** Honey bee, Locomotor activity, Light, Wavelength

## Abstract

Light plays a crucial role in honey bee (*Apis mellifera*) behavior by influencing foraging, navigation, and locomotor activity (LMA). While the effects of light on LMA have been previously documented, the specific roles of different wavelengths remain unknown. In this study, we investigated how exposure to specific infrared (IR, 849 nm), green (528 nm), blue (447 nm), and ultraviolet (UV, 372 nm) wavelengths, as well as their combinations, affects LMA. Specifically, using a custom-built illumination setup and the Api-TRACE video tracking system, we monitored and analyzed bee movement in a homogeneously illuminated environment. Our analysis revealed significant differences in LMA depending on the wavelength to which the bees were exposed. This study demonstrated that the green light promoted LMA. On the other hand, UV light suppressed the LMA of honey bees. The suppression was even greater when the UV light was combined with the blue light. That information can be applied to experimental standardization, the design of flight-room environments, and the management of colonies under artificial illumination.

## INTRODUCTION

Color vision enables bees to recognize flowers, navigate, and forage for food (de Ibarra et al., 2014; Vasas et al., 2019). They possess three types of photoreceptors in their retina, with absorption peaks at 344 nm (UV or short type), 436 nm (blue or medium type), and 544 nm (green or long type) (Menzel & Blakers, 1976; Peitsch et al., 1992; Aurore et al., 2012). Color discrimination is based on an opponency system (Kien & Menzel, 1977; Riehle, 1981; Yang et al., 2004).

The light perception system begins in the retina, and the first stage of processing occurs in the lamina. Color-opponent neurons appear in the medulla and lobula (Yang et al., 2004; Paulk et al., 2009). Then, the lobula sends information for higher-order processing to the anterior optic tube (AOTu) (Paulk et al., 2009; Mota et al., 2011). The AOTu receives chromatic input from the lobula and processes UV, blue, and green light separately in distinct subunits. Green light dominantly activates both the dorsal and ventral lobes, blue light activates the dorsal lobe, and UV light activates the ventral lobe (Mota et al., 2013). The AOTu sends projections to higher-level brain centers such as the lateral protocerebrum (LP) and the central complex (CC). The LP integrates visual information, with anterior LP neurons being primarily color-sensitive, while the CC regulates movement and orientation based on processed chromatic input (Paulk et al., 2009; Mota et al., 2011; Mota et al., 2013).

Apparently, different wavelengths are utilized for specific tasks. Long wavelengths are used for achromatic tasks, such as motion detection (Lehrer et al., 1988), edge detection (Lehrer et al., 1990; Vasas et al., 2017), as well as for chromatic vision. Medium and short wavelengths contribute to chromatic vision (Aurore et al., 2017). Short wavelengths play a particular role in polarized light perception and, in turn, in navigation (Menzel & Blakers, 1976; Peitsch et al., 1992).

Locomotor activity (LMA) measurements indicate circadian rhythm patterns (Erdem et al., 2023). Circadian rhythm is essential for the efficiency of colony tasks, particularly in securing large amounts of food through foraging (Bloch et al., 2017). In honey bees factors such as age, division of labor (Beer & Helfrich-Förster, 2020; Bloch et al., 2001), social environment (Meshi & Bloch, 2007; Bloch, 2010; Eban-Rothschild et al., 2012; Beer et al., 2016; Fuchikawa et al., 2016; Siehler et al., 2021), exposure to chemicals (Tackenberg et al., 2020; Arslan et al., 2023; Erdem et al., 2023), and temperature variations (Moore & Rankin, 1993; Giannoni-Guzmán et al., 2021) influence LMA. Among the factors, light is a strong environmental component that affects LMA (Moore & Rankin, 1993; Spangler, 1973; Erdem et al., 2023; Kim et al., 2024).

Regarding light, there are many studies focusing on the effect of different wavelengths on the activities of several animal species. In the fruit fly (*Drosophila melanogaster*), UV light increased activity compared to darkness (Sakai et al., 2002). Additionally, the activity was higher under blue and green light than red light (Subramanian et al., 2009). In German cockroaches (*Blattella germanica*), UV light elicited higher activity than green light in a 12-hour light and 12-hour dark regimen. Meanwhile, green light caused higher activity than UV in 24-hour constant light conditions (Leppla et al., 1989). In a related study, 30-minute exposure to green light had a stimulating effect; however, UV was observed to increase immobility in American cockroaches (*Periplaneta americana*) (Zhukovskaya et al., 2017). Two notable studies on movement-light interaction of insects showed that the average walking speed for the parasitoid wasp (*Aphidus ervi*) was ranked as UV, green, blue, and red (Cochard et al., 2017), while for the flower bug (*Orius sauteri*), the order was the reverse (Wang et al., 2013). Zebra fish (*Danio rerio*) displayed higher activity between purple to orange wavelength ranges (415 – 595 nm) than red and IR (Di Rosa et al., 2015; Waalkes et al., 2024). In rodents, mean hourly activity count during photophase was in the order of red, green, and blue in the nocturnal Namaqua rock mouse (*Micaelamys namaquensis*); however, the order was red, blue, and green for the diurnal four-striped field mouse (*Rhabdomys pumilio*) (Van Der Merwe et al., 2019). Thus, different wavelengths of light have various effects on animal species. Therefore, we hypothesize that exposure to different wavelengths of light will cause distinct, observable variations in honey bee locomotor activity. This hypothesis will be supported if significant variations in LMA are observed in bees exposed to different wavelengths of light.

Standardization is necessary for LMA and circadian rhythm experiments. Some experiments were conducted using fluorescent light sources (Spangler, 1973) that emit very low levels of UV (FDA, 2017). In other experiments, light-emitting diodes (LEDs) were used, which do not emit UV light (Kim et al., 2024; Erdem et al., 2023). This lack of standardization is equally prevalent in bee flight rooms, where illumination has varied from solely fluorescent light (Jay, 1964; Poppy & Williams, 1999) or LEDs (Buatois et al., 2024) to setups Supplementaryed with additional UV and blue light (van Praagh & Velthuis, 1971), only UV (Czoppelt et al., 1980; Pernal & Currie, 2001), only blue light (Kefuss, 1978), or blue-tinted tracing cloth (Nye, 1962), all without a standardized wavelength protocol. In particular, there had to be standardization in critical experiments, such as on brood development (Czoppelt et al., 1980; Kefuss, 1978), and foraging (Poppy & Williams, 1999; Buatois et al., 2024). We believe this issue remains a fundamental caveat for illumination and argue the imperative for its standardization in future research on LMA, circadian rhythms, foraging behavior, and honey bee colony management.

## RESULTS

We analyzed LMA results separately for the 24-hour period and the last 12 hours. We used the last 12 hours to examine the bees’ LMA after waking (Supplementary Figure 1) and to exclude the low activity during sleep, which was seen in all groups.

First, we statistically evaluate the LMA for the 24 h report the results in Supplementary Table S1. According to the Shapiro-Wilk test, the data for all groups deviated significantly from a normal distribution (p <.05). According to the Shapiro-Wilk test, all groups did not follow the normal distributions (*p* < 0.05). To address this, we used the nonparametric Kruskal-Wallis test to assess differences in LMA across groups. Kruskal-Wallis test indicated the difference across the groups (χ^*2*^(7)□=□23.977, *p*□=□0.001). Then we examined whether there was a difference between the groups using a post-hoc Dunn test. According to the Dunn test, the blue-UV group (to which blue and UV light were applied simultaneously) differed significantly from all other groups (*p* < 0.05). Also, the only-UV group differed from the only-blue, only-green, and blue-green group (the group that received both green and blue light simultaneously) (*p* < 0.05) (Figure 2). There was no difference between all other groups (*p* > 0.05). The results of the Dunn test comparing the groups are given in detail in Supplementary Table S2. Additionally, both the means and medians of the LMA of the only-UV and the combination of UV with other lights were lower than in the darkness treatment (IR group). Meanwhile, the blue and green combination, blue-only, and green-only groups’ means and medians were higher than the darkness (Supplementary Table S1). It was clearly seen that LMA bottomed out when UV and blue light were combined.

**Figure 1.**
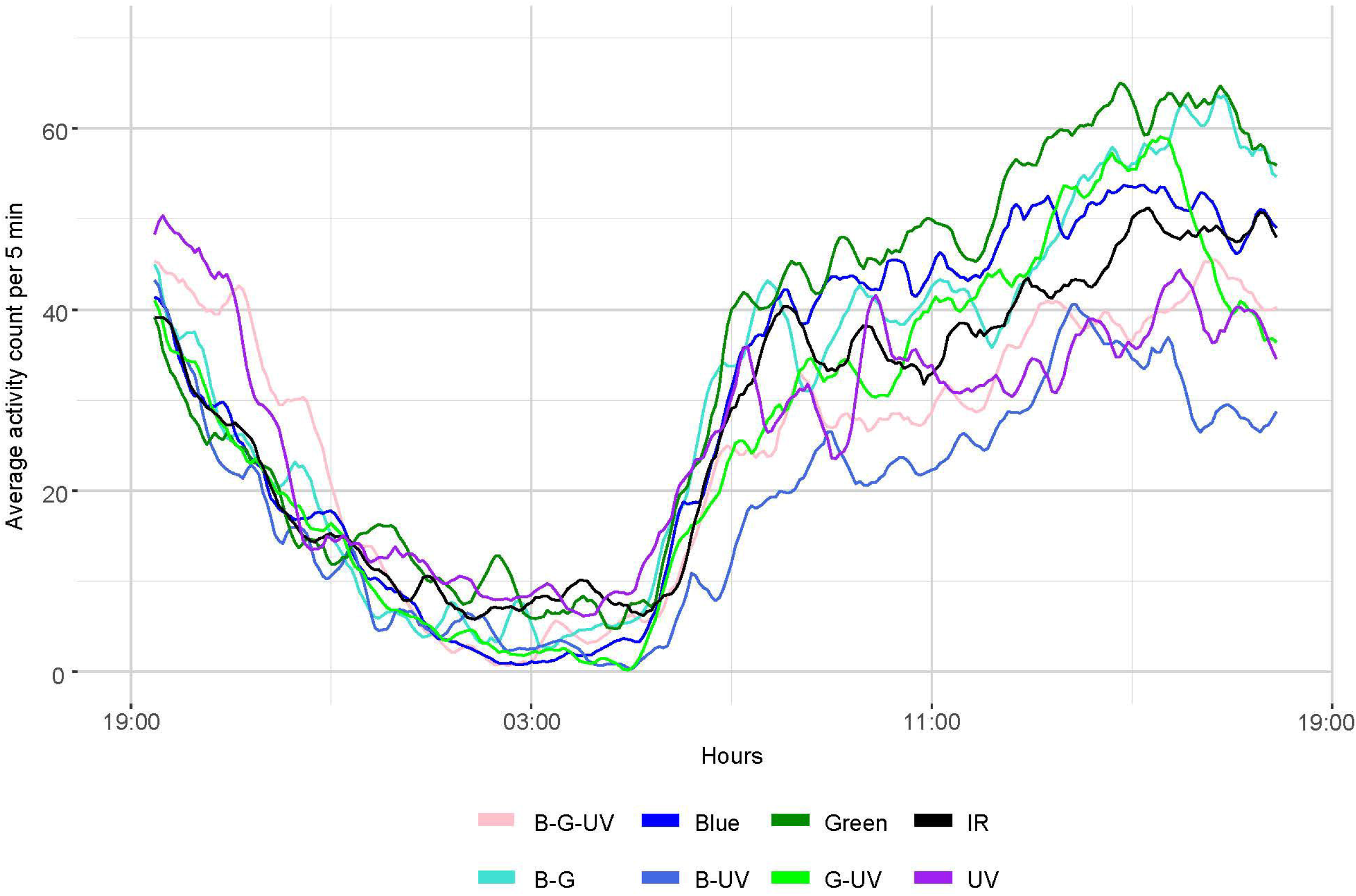
The plot shows changes in daily locomotor activity. B-G-UV is the combination of blue, green, and UV; B-G is blue and green; B-UV is blue and UV; and G-UV is green and UV. A moving-average filter was applied when plotting the graph, accepting 6 data points as input at a time, to calculate an average LMA for each 30-minute interval.

**Figure 2.**
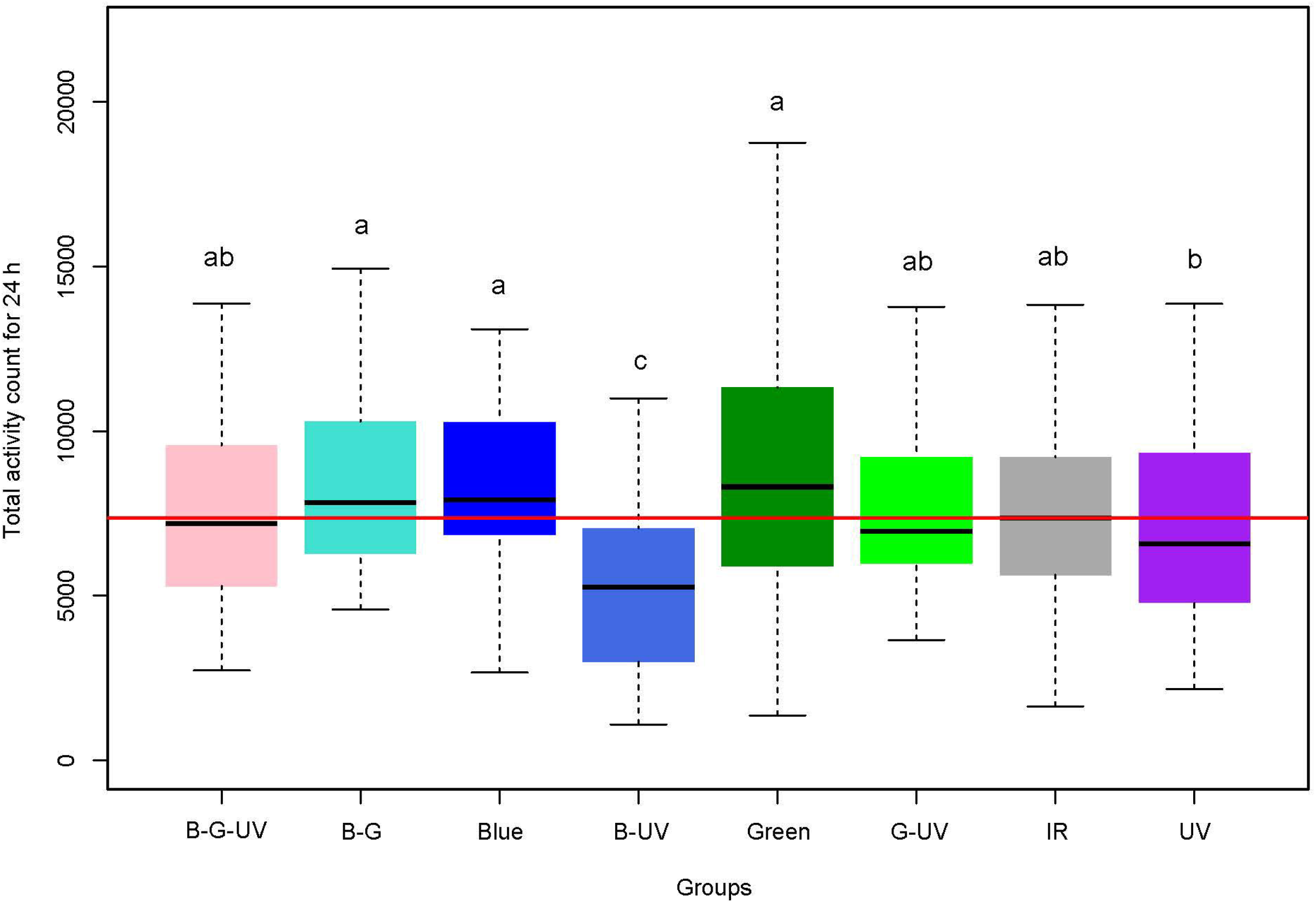
The box plot illustrates comparisons of LMA in response to the light of different wavelengths for 24 hours; B-G-UV is the combination of blue, green, and UV; B-G is blue and green; B-UV is blue and UV; and G-UV is green and UV. When the same letters appear on the bars, it indicates that there is no statistically significant difference between the groups according to Dunn’s test. The data are presented as the median of the total activity count, along with quantiles. The red line indicates the median of the IR group.

Secondly, the LMA in the last 12 hours of the experiment was analyzed statistically, and the basic statistics are presented in Supplementary Table S3. Because the majority of the groups (except blue-green, blue, and green) follow normal distributions according to the Shapiro-Wilk test (*p* > 0.05), a one-way ANOVA test was used to compare the groups’ LMA. The results indicated a significant difference among the groups’ LMA (*F* (7, 239) = 6.21, *p* < 0.001). Post-hoc comparisons using Tukey’s HSD test (Figure 3, Supplementary Table S4) revealed that the only-green illumination exhibited significantly higher LMA than the blue-green-UV group (which received blue, green, and UV wavelengths simultaneously), blue-UV, and only-UV groups (*p* < 0.05). Both blue-green and only-blue also showed significantly higher activity than blue-UV (*p* < 0.05). Additionally, green-UV differed slightly from the blue-UV group (*p* = 0.048). No other pairwise differences reached statistical significance. These results suggest that blue-UV consistently showed the lowest activity, while only-green showed the highest.

**Figure 3.**
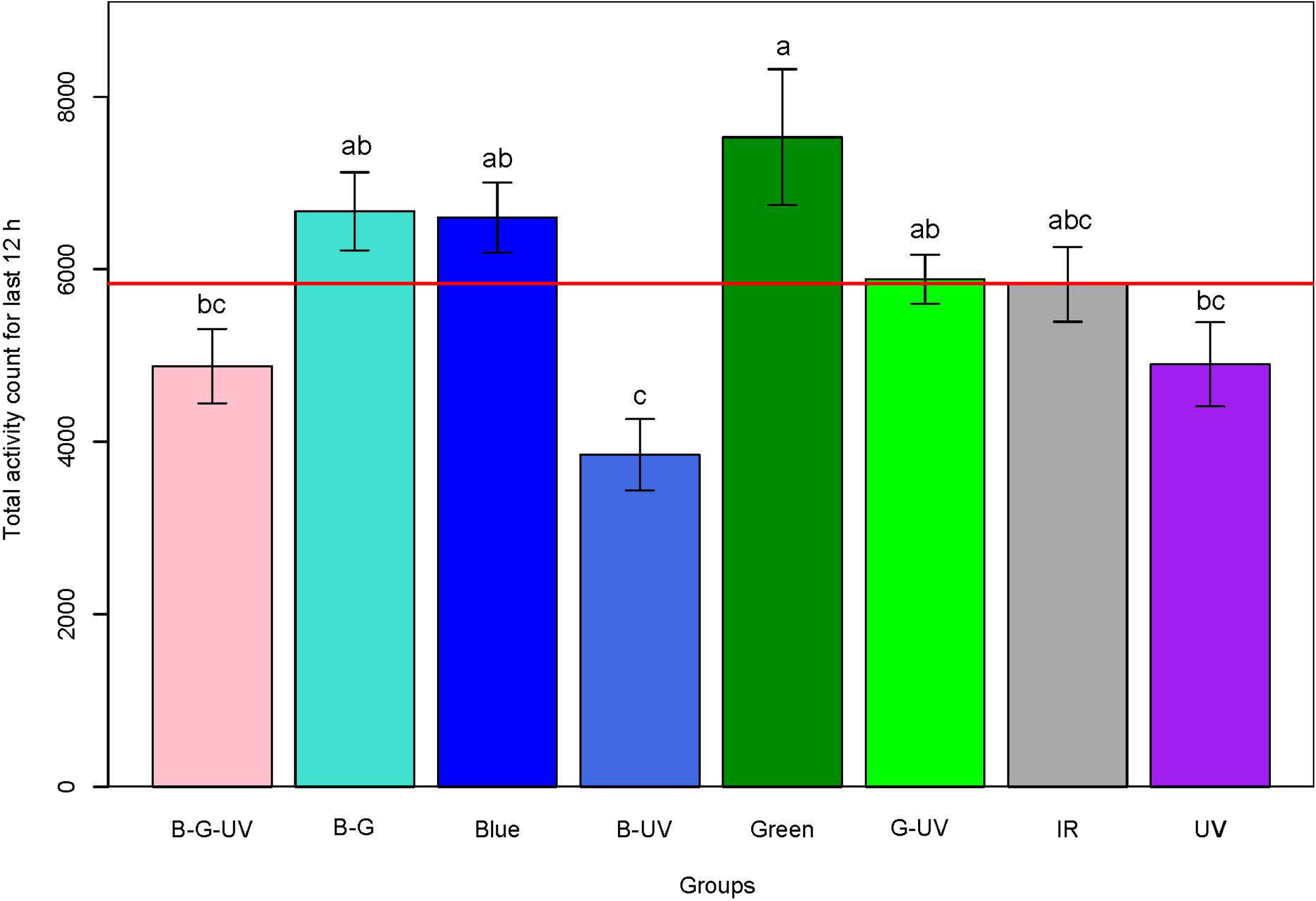
The box plot illustrates comparisons of LMA in response to the application of light of different wavelengths for the last 12 hours; B-G-UV is the combination of blue, green, and UV; B-G is blue and green; B-UV is blue and UV; and G-UV is green and UV. When the same letters appear on the bars, it indicates that there is no statistically significant difference between the groups according to the Tukey test. The data is presented as the mean of the total activity count with ± standard error. The red line indicates the mean of the IR group.

All results of our analysis confirm that the LMA of honey bees is significantly modulated by light wavelength. Specifically, bees exposed to green light showed the highest levels of activity, significantly exceeding those exposed to combinations containing UV light. The combination of blue and UV light yielded the lowest LMA values, indicating a suppressive effect. These findings support the hypothesis that distinct wavelengths differentially influence bee motor behavior. The pronounced activity suppression in UV-containing conditions, particularly blue–UV and green– blue–UV, suggests that UV light plays a dominant inhibitory role on LMA.

## DISCUSSION

Our study demonstrated that different wavelengths of light and combinations significantly affect the LMA of honey bees. Notably, exposure to UV light resulted in a substantial reduction in LMA. When combined with blue light, UV exposure led to the lowest activity levels observed, even lower than those recorded in the IR (darkness) condition. Whereas, green light elevates the LMA of honey bees.

In our study, although the LMA averages of the green, blue, and green-blue light groups were higher than those in the dark, no significant difference was observed across these light treatments. According to previous research, the irradiance values of 4.317 μW/cm^2^ for the 350 nm LED, 3.466 μW/cm^2^ for the 435 nm LED, and 2.668 μW/cm^2^ for the 565 nm were enough to stimulate photoreceptors (Mota et al., 2013; calculations for transformation of photon counts/cm2/s to μW/cm^2^ given in Supplementarys). In our experiments, the irradiance value was 12 μW/cm^2^ for each group. Thus, this value is sufficient to stimulate the photoreceptors. However, this value may not be sufficient to fully saturate the photoreceptors or to produce a very strong stimulus in the relevant visual regions of the brain.

The decrease in LMA under UV and UV-blue light may be explained by neural pathways in the bees. First, it has been shown that UV can have excitatory or inhibitory effects on certain neurons in the lobula (Yang et al., 2004). Also, certain neurons were shown to be affected by both blue and UV light (Yang et al., 2004). Considering that the combination of UV and blue light has the greatest activity-reducing effect, the dual inhibitory neurons (UV-/B-) and dual excitatory opponent (UV+/B+/G^−^) neurons could be responsible for the observed effect on LMA. In addition, it is shown that definite cross-sectional areas in AOTu are activated by both blue and UV light stimulation (Mota et al., 2013).

The effect of wavelengths on activity should be unique for each organism. In our study, maximum activity was determined under green light conditions. However, in other organisms (Subramanian et al., 2009; Cochard et al., 2017; Wang et al., 2013; Waalkes et al., 2024; Van Der Merwe et al., 2019), activity under green light was not at the top. Besides, as mentioned above, animal species also vary in their response to UV light in their activities (Cochard et al., 2017; Leppla et al., 1989; Zhukovskaya et al., 2017). In our study, we observed elevation under UV light at the beginning of our experiment (Figure 1). Thus, UV light initially acted as an excitatory stimulus before its suppressive effects took hold. From a behavioral perspective, it has been reported that UV light is used primarily for navigation (Menzel & Blakers, 1976; Peitsch et al., 1992). It has also been found that bees exhibit a stronger attraction towards UV light than other wavelengths (Nouvian & Galizia, 2020). When bees were suddenly confronted with unpolarized, directionless, and homogenized UV light, they might have become confused and exhibited rapid searching behavior.

In conclusion, our study provides strong evidence that UV light plays a critical role in modulating honey bee locomotor activity. That likely occurs through the specific activation of neural pathways involved in visual processing. The significant suppression of LMA under UV and UV-blue light conditions suggests an adaptive response that may help bees regulate their activity levels in natural environments, such as fluctuations in foraging activity throughout the day that UV light reaches highest level at the middle of day (2052 μW/cm^2^) and lowest at afternoon (267 μW/cm^2^) at the middle latitudes (Liu & Zhang, 2015). Further research should aim to elucidate the underlying neural mechanisms and explore the broader ecological and agricultural implications of these findings.

## MATERIALS AND METHODS

The bees used were obtained from two colonies located on the Middle East Technical University campus. Returning forager bees were captured by placing plastic wire mesh over the hive entrances, and the bees that gathered on it were collected into small cages.

A custom-made illumination setup was built for the experiments (see the Supplements for details; Supplementary Figure S1).). We utilized light-emitting diodes (LEDs) with peak wavelengths of 528 nm (green), 447 nm (blue), 372 nm (UV), and 849 nm (IR), which were verified using a spectrometer (see Supplements for details, Supplementary Figure S2). These wavelengths fell within the established spectral sensitivity intervals of the honey bee’s color receptors (Supplementary Figure S3). The LEDs were connected to adjustable DC power supplies, whose current and voltage were regulated to deliver a consistent irradiance of 12 μW/cm^2^ for each wavelength. Irradiance values were measured just before the experiment using the TSL2561 light sensor breakout card connected to an Arduino circuit (the code, circuit diagram, and details on irradiance conversion were provided in the GitHub repository at github.com/baburerdem/IrradianceSensor).

Bees were placed in the glass tubes (^Ø^16 x 100 mm) with a 2 mm-diameter hole in the middle to allow ventilation. The mouth of each tube was covered with a lid containing fondant sugar for the bees to feed on, and cheesecloth to prevent them from sticking to the sugar. The tubes were placed horizontally, side by side, inside the climate chamber set to 32 °C and 60 % relative humidity. A webcam (Logitech C920; the IR filter was removed) was placed inside the chamber, parallel to the tubes and at a distance that covered all tubes (Supplementary Figure S1). Eight separate experiments were conducted with only green, blue, UV, and combinations of green-blue, green-UV, blue-UV, green-blue-UV, and only IR (dark for bees) LEDs on. The bees were replaced at the beginning of each experiment, and 32 were used in each. The experiments started at 7:00 pm and lasted 24 hours. Dead bees were identified and removed from the data; the last sample sizes are given in Supplementary Table S1.

Previous studies used a commercial device to measure LMA, which counted bee movements in the tubes using IR sensors (Giannoni-Guzmán et al., 2014; Beer et al., 2016; Tackenberg et al., 2020; Arslan et al., 2023; Erdem et al., 2023). In this device, the tubes were positioned vertically on top of each other so that the light could not reach the tubes homogeneously. In our experiments, we aligned the tubes horizontally and used a computer vision (CV) algorithm we had previously developed (Erdem et al., 2024). The system called Api-TRACE: Honey Bee Tracking in Constrained Environments was a computer vision-aided system to track honey bees in a constrained environment (Erdem et al., 2024). Using this system, we measured LMA by tracking the movements of the bees in horizontally arranged, homogeneously lighted tubes.

24-hour video recordings were processed with the Api-TRACE Video Processing Module (github.com/baburerdem/Api-TRACE). Activity counts were extracted using the code, similar to methods of counting the passage of bees across the midpoint of the tubes (Rosato & Kyriacou, 2006; Giannoni-Guzmán et al., 2014), and were used in the statistical analysis.

## ACKNOWLEDGMENTS

We thank Prof. Dr. Alpan Bek from the Physics Department at Middle East Technical University for his help in ensuring precise wavelength measurements of the LEDs used in the experiments, and Prof. Dr. Tugrul Giray from the Department of Biology at the University of Puerto Rico for his feedback and for revising the draft. We thank Middle East Technical University [BAP grant number: ADEP-302-2024-11468]; The Scientific and Technological Research Council of Türkiye [T^ÜBİ^TAK grant number: 122E014] for supporting this work.

## DATA AVAILABILITY

The data are available in the Zenodo repository: https://doi.org/10.5281/zenodo.17414161

## AUTHOR CONTRIBUTIONS

Babur Erdem: Conceptualization, Methodology, Software, Formal analysis, Investigation, Writing - Original Draft, Visualization. Iremnur Fidan: Investigation. Ali Emre Turgut: Methodology, Resources, Funding acquisition. Erol Sahin: Methodology, Resources. Ayse Gul Gozen: Writing - Original Draft. Hande Alemdar: Methodology, Resources, Writing - Original Draft, Funding acquisition.

