## Supplementary for "Specific Wavelengths of Light Modulate Honey Bee Locomotor Activity"

**SUPPLEMENTS**

Materials and methods

- Illumination and experiment setup

To obtain green, blue, and UV lights, two power LEDs each were used. Each pair of LEDs was connected in parallel. There were eight parallel connected mini LEDs to provide IR illumination. The LEDs were placed by applying thermal paste and screwing them onto a metal plate. Two mini fans were installed beneath the metal plate to prevent the LEDs from overheating. The metal plate was attached to the aluminum profiles with the LEDs facing upwards. Thanks to the LEDs facing upwards, the light did not hit the glass tubes directly, thus providing homogeneous illumination.

The illumination setup was placed on the upper shelf of the climate cabin. A webcam was attached to the bottom of the upper shelf. On the lower shelf, glass tubes were lined up on a white background, held in place with double-sided tape (Figure S1).

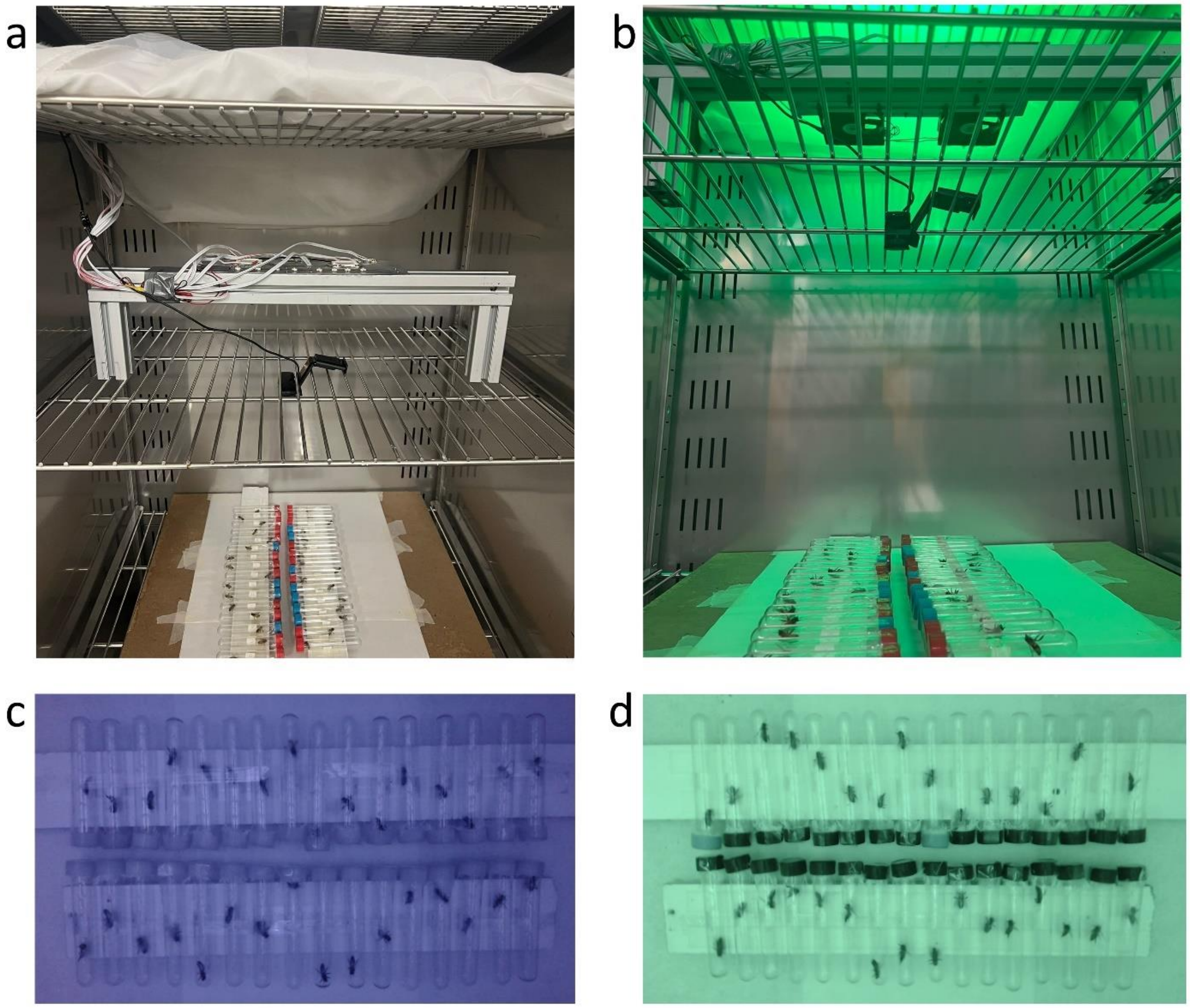


Figure S1. View of the experimental setup from different angles with the illumination setup off (a) and with the illumination setup on (b). Camera view under IR (c) and green light (d).

- Wavelength measurement of the LEDs

The spectrometer (Andor SR750) was connected to the computer with Andor Solis software. A 532 nm laser source at 0.01 W was used for calibration. The parameters were set to an exposure time of 10s, accumulation number of 10, EM gain active, and readout speed of 50 kHz at 16 bits. Measurements were taken and recorded for each LED. Plots were drawn from the obtained data using Origin software. The 530.5 nm reference measurement was taken from the 532 nm laser, which was the reference source, with a calibration error of 1.5 nm.

Peak points of the UV, blue, green, and IR LEDs were at 372 nm, 447 nm, 528 nm, and 849 nm, respectively (Figure S2).

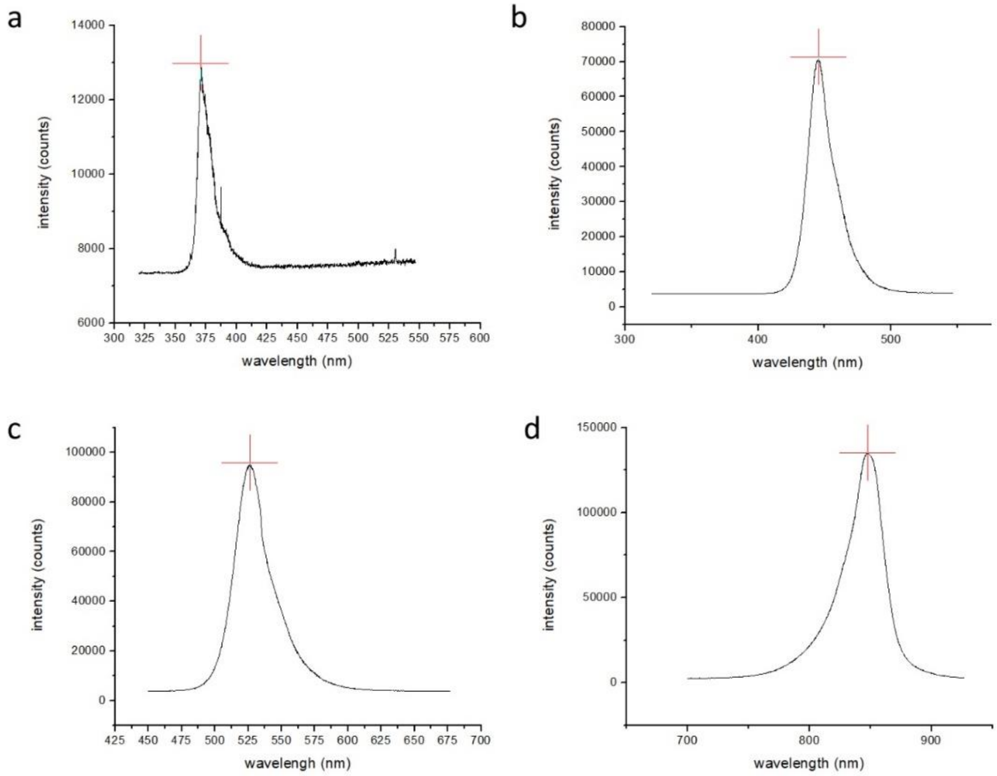


Figure S2. Peaks of the wavelengths of the LEDs. The peak point of the UV LED is 372 nm (a), blue is 447 nm (b), green is 528 nm (c), and IR is 849 nm (d).

- Comparison of the wavelength interval of the LEDs and the spectral sensitivity of the color receptors of honey bees.

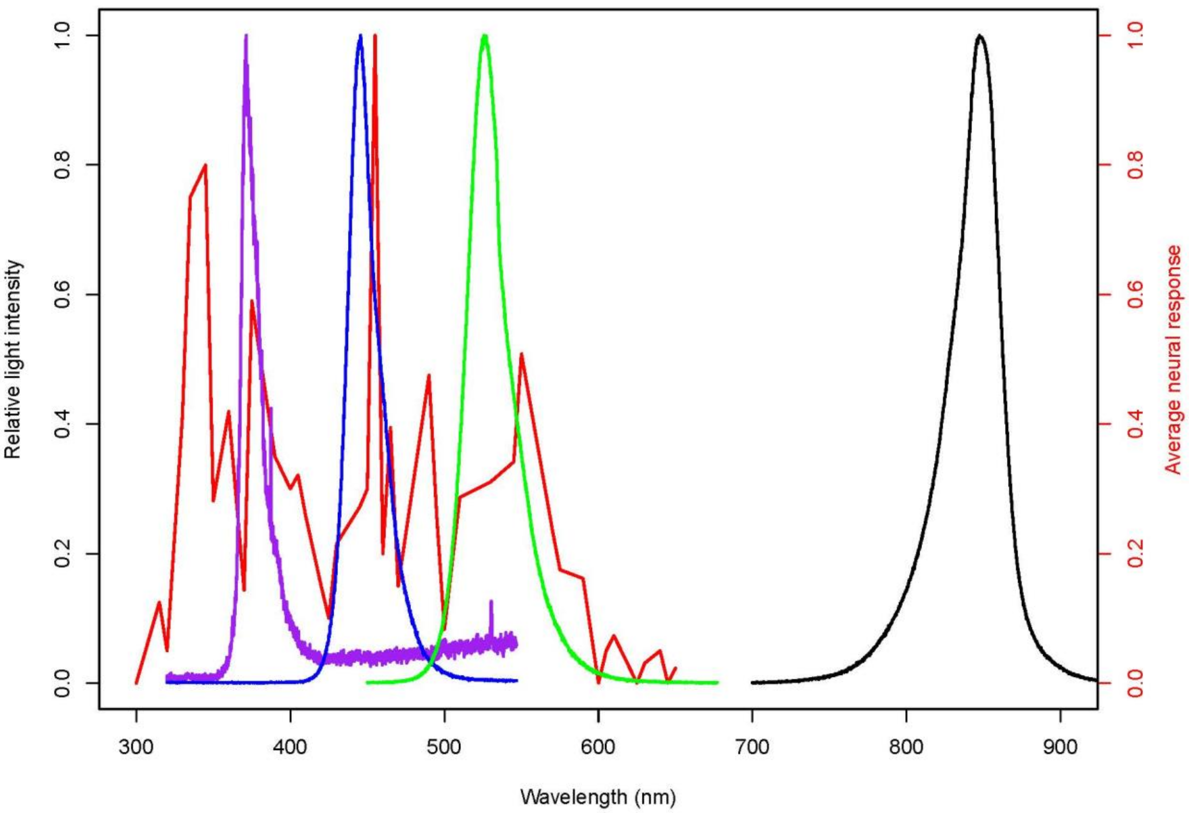


Figure S3. Relative light intensities of our UV (purple), blue (blue), green (green), and IR (black) LEDs, and the average neural response of the color neurons (red) are illustrated. In the plot, it is seen that the wavelength range in which UV, blue, and green LEDs emit photons coincides with the spectral sensitivity range of color neurons. The range of the IR LED falls outside the spectrum to which neurons are sensitive. For each LED, the smallest reading in the raw data received from the device was subtracted from all values to reduce the effect of background light. The values were then divided by the largest value to obtain the relative light intensity curves. The average neural response curve is the mean of the data obtained from 22 neurons, as reported by Menzel and Blakers (1976). This data was obtained from the appendix of the article by Vasas et al. (2019).

- Calculation of the adequacy of the irradiance value we applied in the experiment to stimulate the photoreceptors.

Mota et al. (2013) used LEDs at 350, 435, and 565nm and a light flux of 3.8 × $10^6$ photon counts/$cm^2$/s for stimulation and 7.6 x $10^6$ photon counts/$cm^2$/s for maximum stimulation of photoreceptors.

These light fluxes are converted to the unit of measurement we use, μW/cm², as follows:
First, find the energy of a single photon of different wavelengths.
The energy of a single photon is given by:

$$E_{\text{photon}} = \frac{hc}{\lambda}$$

Where:

$h$ = 6.626 x $10^{-34}$ J·s (Planck's constant), $c$ = 3.0 x $10^8$ m/s (speed of light), $\lambda$ is the wavelength in meters.

Calculating for each wavelength:

For 350 nm:

$$E_{photon} = \frac{(6.626 \times 10^{-34})(3.0 \times 10^{8})}{350 \times 10^{-9}}$$

$$E_{\text{photon}} = 5.68 \times 10^{-19} \text{ J}$$

For 435 nm:

$$E_{photon} = \frac{(6.626 \times 10^{-34})(3.0 \times 10^{8})}{435 \times 10^{-9}}$$

$$E_{\text{photon}} = 4.56 \times 10^{-19} \text{ J}$$

For 565 nm:

$$E_{photon} = \frac{(6.626 \times 10^{-34})(3.0 \times 10^{8})}{565 \times 10^{-9}}$$

$$E_{\text{photon}} = 3.51 \times 10^{-19} \text{ J}$$

Secondly, compute the power density (P).

$$P = (\text{Photon Flux}) \times E_{\text{photon}}$$

Where:

$$\text{Photon Flux} = 8 \times 10^{6}\ \text{photons/cm}^{2}/\text{s}$$

For 350 nm:

$$P = (7.6 \times 10^{6}) \times (5.68 \times 10^{-19})$$
$$P = 4.317 \times 10^{-12}\ \ W/cm^{2}$$

For 435 nm:

$$P = (7.6 \times 10^{6}) \times (4.56 \times 10^{-19})$$
$$P = 3.466 \times 10^{-12}\ \ W/cm^{2}$$

For 565 nm:

$$P = (7.6 \times 10^{6}) \times (3.51 \times 10^{-19})$$
$$P = 2.668 \times 10^{-12}\ \ W/cm^{2}$$

Lastly, convert $W/cm^2$ to $\mu W/cm^2$.

Since 1 W = $10^6$ μW, multiply by $10^6$:

For 350 nm:

$$4.544 \times 10^{-12} \times 10^{6} = 4.317\ \mu\text{W/cm}^{2}$$

For 435 nm:

$$3.648 \times 10^{-12} \times 10^{6} = 3.466\ \mu\text{W/cm}^{2}$$

For 565 nm:

$$2.808 \times 10^{-12} \times 10^{6} = 2.668\ \mu\text{W/cm}^{2}$$

Thus, the irradiance values of 4.317 $\mu W/cm^2$ for the 350 nm LED, 3.466 $\mu W/cm^2$ for the 435 nm LED, and 2.668 $\mu W/cm^2$ for the 565 nm LED are sufficient to stimulate the photoreceptors at the maximum level. According to this result, the irradiance of 12 $\mu W/cm^2$ we applied at each wavelength should be sufficient to stimulate the photoreceptors.

- Results

Table S1. Basic statistics for 24-hour comparison.

| | n | Mean | Std. Dev. | Median | Std. Err. | First Quantile | Third Quantile |
|---|---|---|---|---|---|---|---|
| B-G-UV | 32 | 7279.281 | 2848.928 | 7199.500 | 503.624 | 5325.250 | 9461.000 |
| B-G | 31 | 8833.065 | 3352.877 | 7824.000 | 602.194 | 6284.000 | 10286.500 |
| B-UV | 31 | 5474.742 | 3442.056 | 5259.000 | 618.212 | 2991.500 | 7048.000 |
| Blue | 32 | 8497.469 | 2928.480 | 7919.000 | 517.687 | 6916.750 | 10248.500 |
| G-UV | 32 | 7710.469 | 2421.376 | 6954.000 | 428.043 | 5986.250 | 9002.250 |
| Green | 29 | 9772.793 | 6959.965 | 8308.000 | 1292.433 | 5902.000 | 11323.000 |
| IR | 31 | 7980.903 | 4010.250 | 7359.000 | 720.262 | 5632.500 | 9206.000 |
| UV | 29 | 7675.690 | 4262.023 | 6581.000 | 791.438 | 4788.000 | 9329.000 |

Table S2. Dunn test result matrix with p-values for 24-hour comparison.

| | B-G-UV | B-G | Blue | B-UV | Green | G-UV | IR |
|---|---|---|---|---|---|---|---|
| B-G | 0.050 | | | | | | |
| Blue | 0.054 | 0.478 | | | | | |
| B-UV | 0.011 * | < 0.001 *** | < 0.001 *** | | | | |
| Green | 0.060 | 0.474 | 0.496 | < 0.001 *** | | | |
| G-UV | 0.278 | 0.144 | 0.155 | 0.002 ** | 0.164 | | |
| IR | 0.332 | 0.115 | 0.124 | 0.003 ** | 0.132 | 0.440 | |
| UV | 0.420 | 0.036 * | 0.039 * | 0.020 * | 0.044 * | 0.219 | 0.267 |

Table S3. Basic statistics for the last 12-hour comparison.

| | n | Mean | Std. Dev. | Median | Std. Err. | First Quantile | Third Quantile |
|---|---|---|---|---|---|---|---|
| B-G-UV | 32 | 4875.313 | 2433.231 | 4978.000 | 430.139 | 3080.500 | 6713.000 |
| B-G | 31 | 6670.000 | 2516.529 | 6394.000 | 451.982 | 4871.500 | 7890.500 |
| B-UV | 31 | 3848.323 | 2298.055 | 3391.000 | 412.743 | 2137.000 | 5057.500 |
| Blue | 32 | 6598.375 | 2285.326 | 6420.000 | 403.992 | 5695.750 | 7135.250 |
| G-UV | 32 | 5881.031 | 1598.159 | 5866.500 | 282.517 | 4992.000 | 6874.750 |
| Green | 29 | 7529.069 | 4246.230 | 6100.000 | 788.505 | 4595.000 | 9165.000 |
| IR | 31 | 5824.129 | 2413.125 | 5285.000 | 433.410 | 4248.000 | 6986.000 |
| UV | 29 | 4896.414 | 2631.732 | 4634.000 | 488.700 | 2993.000 | 6748.000 |

Table S2. Tukey test result matrix with p-values for the last 12-hour comparison.

| | B-G-UV | B-G | Blue | B-UV | Green | G-UV | IR |
|---|---|---|---|---|---|---|---|
| B-G | 0.124 | | | | | | |
| Blue | 0.152 | 1.000 | | | | | |
| B-UV | 0.778 | < 0.001 *** | < 0.001 *** | | | | |
| Green | 0.003 ** | 0.910 | 0.864 | < 0.001 *** | | | |
| G-UV | 0.789 | 0.933 | 0.958 | 0.048 * | 0.223 | | |
| IR | 0.840 | 0.909 | 0.940 | 0.065 | 0.194 | 1.000 | |
| UV | 1.000 | 0.155 | 0.188 | 0.782 | 0.004 ** | 0.826 | 0.871 |
